# Analysing antimicrobial resistance mobility patterns using a diverse dataset of over 8000 bacterial species

**DOI:** 10.64898/2026.03.03.709307

**Authors:** Baofeng Jia, Brian P. Alcock, Amogelang R. Raphenya, Jason Spence, Finlay Maguire, Robert G. Beiko, Andrew G. McArthur, Claire Bertelli, Fiona S.L. Brinkman

## Abstract

2.

Advances in genomics have enhanced surveillance of antimicrobial resistance (AMR), yet the factors governing resistance gene mobility, and therefore their risk of spread, remain poorly characterized. Here, we analyzed AMR gene distributions across thousands of bacterial genomes from NCBI RefSeq, encompassing clinical, agricultural and environmental isolates, to quantify associations with plasmids and predicted mobile genomic islands. AMR genes were identified using the Resistance Gene Identifier, and in addition to plasmid identification, mobile chromosomal elements were predicted using IslandViewer 4. Analysing both the full dataset and subsets of data with corrections for sampling bias, we show that known AMR genes are significantly enriched in mobile regions overall. However, stratification by resistance mechanism supported marked heterogeneity: certain drug classes and mechanisms are strongly associated with mobile elements, whereas others are predominantly chromosomal and non-mobile. Notably, mechanisms with specialized functions showed higher mobility, consistent with their role as “ecological public goods” that need not be present in all cells to confer community-level benefit. Differences were also observed across bacteria with distinct cell envelope structures. Together, these findings lay the groundwork for predictive models of AMR gene mobility and provide a framework for incorporating gene-level mobility into AMR risk assessment and antimicrobial stewardship policy.

**Impact statement:** Understanding which antimicrobial resistance (AMR) genes are most likely to spread is central to managing resistance in clinical, agricultural, and environmental settings. This study reports a large-scale, systematic analysis of antimicrobial resistance (AMR) gene mobility across >8000 bacterial species, spanning diverse clinical and other ecological sources. By linking resistance mechanisms to their genomic location (plasmids, genomic islands, or the rest of chromosomes), our study demonstrates that AMR genes are overall enriched in mobile elements, but that mobility varies substantially by mechanism and bacterial context. In particular, functionally specialized resistance mechanisms are more frequently associated with mobile elements, consistent with an “ecological public goods” model for AMR dissemination. These findings extend beyond gene-specific case studies to identify generalizable patterns that help explain why some resistance determinants spread more readily than others. The breadth of relevance spans clinical microbiology, evolutionary biology, environmental microbiology, agri-foods, and public health. The work provides evidence supporting an improved framework for AMR mobility risk assessment, and antimicrobial stewardship strategies, with direct implications for forecasting the durability of existing and future antibiotics.

**Data summary:** The complete dataset and analysis code used to generate the results can be accessed via the Open Science Framework (OSF) repository under DOI:10.17605/OSF.IO/WE3TX

## 5. Introduction

The discovery of antibiotics in the 1930s, followed by the “golden era” of antibiotic discovery in the 1950s, transformed human and animal health by making many previously lethal infections treatable. Antibiotics now underpin modern medicine, yet the widespread use of these drugs has driven the global emergence of antimicrobial resistance (AMR), including giving rise to multidrug-resistant pathogens, or “superbugs” (1). This threat has intensified in the “post-antibiotic era”, where bacterial evolution outpaces the development of new therapeutics (1). Industrial-scale production and extensive use of antibiotics in humans and animals have created pervasive selective pressure, amplifying and disseminating resistance determinants (2). Recent reviews in 2021 and 2024 highlighted that AMR currently causes an estimated 1.2 million deaths annually. Forecasting suggests an increase to 1.9 million annual deaths attributable to AMR globally by 2050, for a cumulative total of over 39 million deaths by 2050 attributable to AMR (3, 4).

AMR genes are ancient and ubiquitous. *In silico* analyses suggest metallo-β-lactamases emerged ∼1 billion years ago, and serine β-lactamases were mobilized onto plasmids 10–100 million years ago (6,7,8). Resistance genes have also been detected in 30,000-year-old permafrost and caves isolated from human influence for over four million years (9, 10, 11). While some resistance arises from adaptive mutations, many AMR genes are inherently mobile, contributing directly to the spread of clinically relevant resistance and treatment failures (9). For example, *qnr* genes likely originated from *Shewanella* species, and CTX-M β-lactamases from *Kluyvera* species (12, 13). Mobile genetic elements (MGEs) facilitate lateral gene transfer (LGT), enabling AMR dissemination across diverse bacterial species. Plasmids, mobile genomic islands, integrons, transposons, and prophages mediate this transfer - for example NDM-1–harboring plasmids occur in *Providencia, Klebsiella*, and *Acinetobacter*, while *Salmonella* genomic island GI1 carries multiple resistance genes (14, 15). Widespread antibiotic use can drive the stochastic capture of environmental resistance genes by MGEs, which can subsequently be acquired by pathogens.

AMR is a complex global challenge. While stewardship programs mitigate antibiotic abuse and overuse, they are reactive and require sustained, widespread compliance, which remains impractical in developing countries (4, 5). Environmental emergence and rapid dissemination of plasmid-borne *NDM-1* and *MCR-1* have compromised last-resort drugs, carbapenems and colistin, and carbapenemase-producing organisms have increased fivefold since 2014. Despite the threat, major gaps remain in understanding AMR gene mobility, spread, and evolution. Genome sequencing has enabled unprecedented insights, with NCBI RefSeq now containing thousands of high-quality genomes across diverse species, and curated databases such as the Comprehensive Antibiotic Resistance Database (CARD; 16), combined with analytical tools for mobility gene detection (17), allow large-scale studies of AMR dissemination. This includes detection of plasmids, and genomic islands (GI’s; clusters of genes of probable horizontal origin; 18). Here, we present a comprehensive analysis of AMR associations with such MGEs across >39,000 predicted bacterial replicons, representing 8120 named species in NCBI RefSeq. We identify resistance genes associated with mobility and higher risk of global spread across human, other animal, food, and environmental reservoirs. Based on ontology-based analysis of trends, we identify part of an evolutionary framework to predict gene mobility potential, with the long-term goal of informing AMR risk assessment and guiding antimicrobial stewardship policy.

## 6. Section headings

### 6.1 Methods

#### Data collection, storage and overview

The NCBI RefSeq database, containing high-quality, non-redundant, and well-annotated reference genomes, was downloaded on 23 May 2024 (RefSeq release 223) and imported into MicrobeDB (19), a locally maintainable microbial genome database. This dataset comprised 39,089 bacterial accessions, representing 8120 species, 1704 genera, across 45 phyla. Three subsets were derived from the dataset: 1) the complete RefSeq dataset, 2) a randomly selected subset containing up to 300 genomes per genus (for genera <300 genomes, all are included), and 3) a subset containing all genomes from those genera listed in a WHO priority pathogens list for new antibiotic R&D (20), including *Pseudomonas, Salmonella, Klebsiella, Escherichia*, and *Burkholderia*. The overall workflow of the mobility analysis is summarized in Supp Figure 1.

#### Mobile sequence and AMR Profile prediction

Plasmids were identified within each complete RefSeq genome, as annotated. Genomic islands (GIs) were predicted by submitting all chromosomal accessions to IslandViewer 4 (21) using default parameters. The resulting tab-delimited output, which included accession numbers and GI start and stop positions, was used to format a URL syntax to query NCBI E-utilities’ REST API to obtain lists of genes contained within each GI.

AMR genes were predicted using the Resistance Gene Identifier (RGI; 16) v6.0.1 in conjunction with the Comprehensive Antibiotic Resistance Database (CARD; 16) v3.2.9, using DNA FASTA files as input and the parameters ‘--alignment_tool DIAMOND’ and ‘--input_type contig’. RGI output consisted of a tab-delimited table of AMR genes and their genomic coordinates.

#### AMR mobility analysis and statistical tests

For the association analysis, AMR genes predicted by RGI were classified into three categories: 1) non-GI chromosomal regions, 2) genomic islands (predicted by IslandViewer 4), and 3) plasmids. Associations between AMR genes and these categories were evaluated using Fisher’s Exact test, appropriate for categorical data resulting from binary classifications. False discovery rate (FDR) correction was applied using the Benjamini-Hochberg procedure.

For bacterial characteristic associations, AMR genes in non-GI chromosomal regions and mobile elements were further stratified by Gram status (positive or negative), and associations were tested using Fisher’s Exact test. The Gram stain status (diderm versus monoderm) was predicted by and obtained from PSORTdb v4.0 (22). Genomes that had an atypical or indeterminate Gram status prediction were excluded from this analysis.

For continuous variables such as average genome size and local %GC variance, values were plotted against the proportion of AMR genes in mobile elements (plasmids and genomic islands) relative to non-GI chromosomal regions. Higher-order categories, including drug class, resistance mechanism, and generalizability (average number of drug classes to which each gene confers resistance), were determined using the Antibiotic Resistance Ontology (ARO) v3.2.9 within CARD. Observed proportions of AMR genes within non-GI chromosomal regions and mobile elements were visualized using R (v4.1.2), the ggplot2 (23) graphical library, and Excel, highlighting deviations from expected distributions.

For all Fisher’s exact tests, a two by two contingency table was produced to quantify the association between two variables. The test was performed using R (V4.1.2)’s ‘fisher.test()’ function, and the odds ratio (OR) is defined as OR=(a×d)/(b×c) and represents the ratio of the odds of the outcome in one group relative to the other. OR > 1 represents a positive association and OR < 1 represents a negative association.

### 6.2 Results

#### Overview of AMR mobility associations

The dataset comprised 39,089 accessions, including 29,793 chromosomal and 9,295 plasmid sequences. We identified 499,319 AMR determinants among 118,144,860 genes (0.42%). Large-scale mobility analysis revealed that known AMR genes are disproportionately located within mobile genomic regions (p < 0.001, Odds Ratio (OR) 3.01; Fig. 1), particularly plasmids (p < 0.001, OR 0.19). Given the composition of the RefSeq database, certain human-and animal-associated bacterial species are overrepresented; for example, Enterobacteriaceae accounted for 30.1% (11,748) of all accessions (Supp Fig. 2). To address this bias, we performed a dataset reduction (see Methods; Supp. Fig. 3). With this reduced dataset, AMR genes remained disproportionately associated with mobile regions (p < 0.001, OR 2.42).

**Figure 1.**
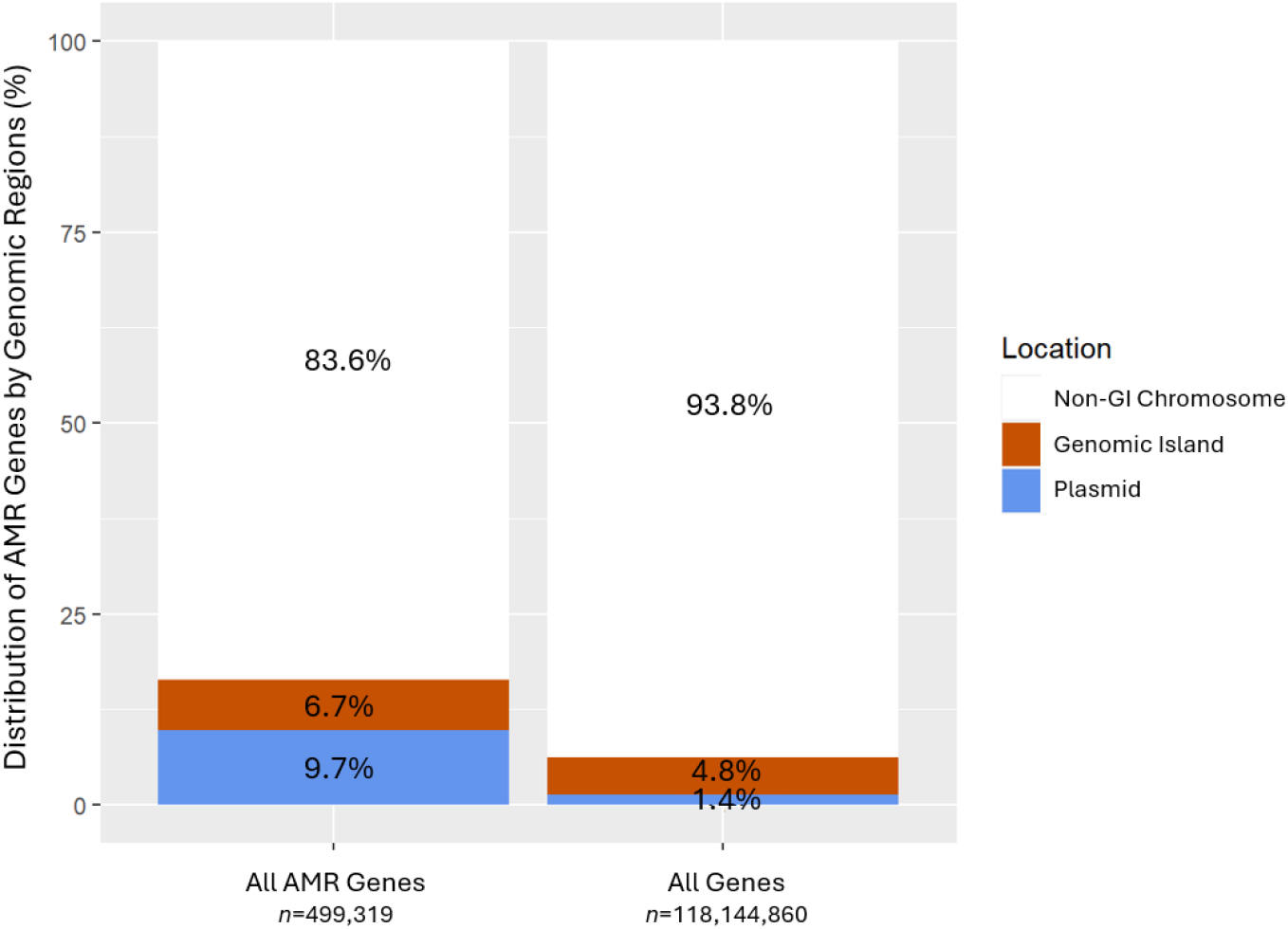
Antimicrobial resistance genes, collectively, are disproportionately found in mobile regions of the genome, particularly plasmids. Accounting for all genes (right bar), the expected proportion of genes (y-axis) on mobile genetic elements was ∼6%. The observed proportion of AMR genes (left bar) was ∼17%. There is a significant association between AMR genes and mobile genetic elements (Fisher’s test, p < 0.001). Of the mobile genetic elements, the majority of the AMR genes are disproportionately found on plasmids (red; p <0.001).

#### Microbial biological and genomic characteristics do not affect an AMR gene’s disproportionate association with mobile elements

To explore whether bacterial traits influence AMR mobility, the dataset was stratified by Gram stain status (diderms versus monoderms) using our approach implemented for updating such predictions for PSORTdb (22), resulting in 27,510 Gram-negative and 11,200 Gram-positive accessions. AMR genes were disproportionately associated with mobile elements in both groups (Supp. Fig. 4). A minor but significant difference was observed: Gram-negative bacteria were slightly more likely to harbor mobile AMR genes than Gram-positive bacteria (p < 0.001, OR 1.12), and AMR genes were more localized to genomic islands in Gram-positive species than Gram-negative species. Other genomic characteristics, including average genome size, local %GC variance, and the antibiotic drug classes targeted by AMR genes, did not alter this association.

#### AMR gene mobility is associated with AMR mechanism of action

Using the Antibiotic Resistance Ontology (ARO), we classified AMR genes into seven resistance mechanism categories: antibiotic efflux (n = 292,508), reduced permeability to antibiotics (= 15,268), antibiotic target alteration (n = 133,398), antibiotic target protection (n = 8,691), antibiotic inactivation (n = 68,340), and antibiotic target replacement (n = 13,404). Distinct mobility trends emerged (Fig. 2).

**Figure 2.**
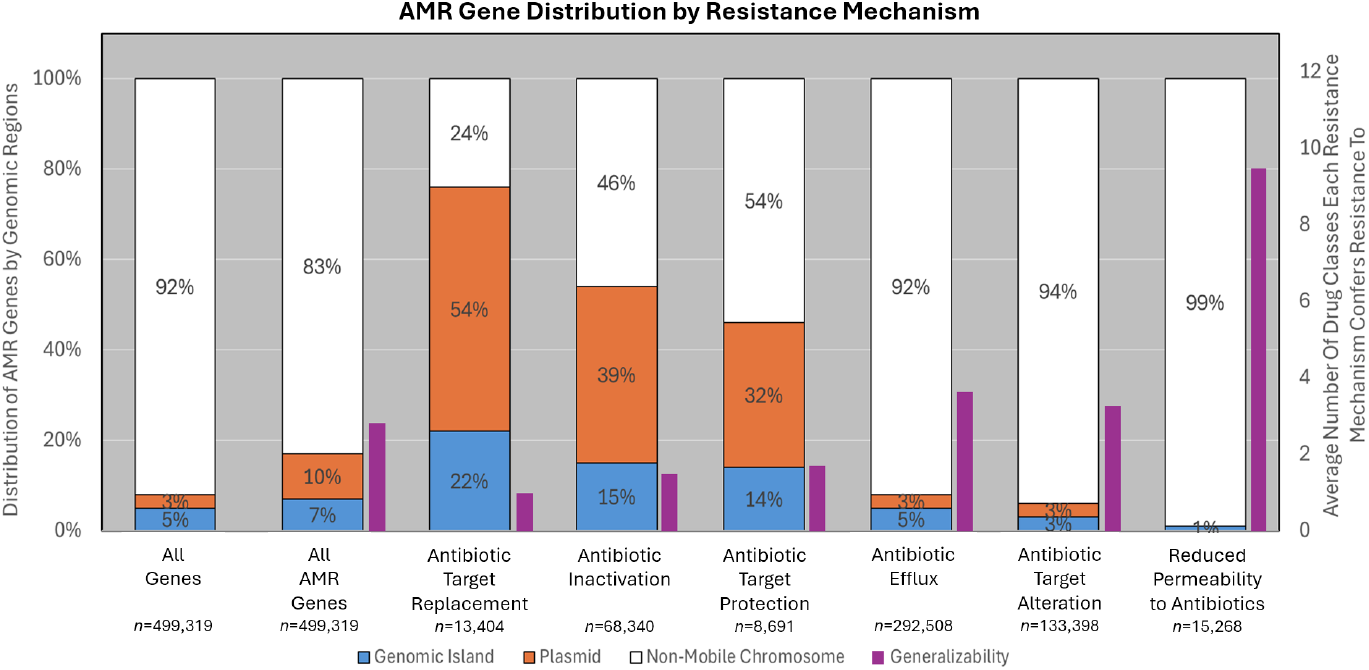
The resistance mechanism of antimicrobial resistance genes correlates with AMR gene distribution. Those genes with low generalizability have higher fitness costs and are associated with mobile elements. All AMR genes are classified into 6 categories: 1) Antibiotic Target Replacement, 2) Antibiotic Inactivation, 3) Antibiotic Target Protection, 4) Antibiotic Efflux, 5) Antibiotic Target Alteration, 6) Reduced Permeability to Antibiotics. A mobility trend can be seen here with reduced permeability to antibiotic (p < 0.001), antibiotic efflux (p < 0.001) and target alteration (p < 0.001) being non-mobile chromosome associated (white) while target protection (p < 0.001), target inactivation (p < 0.001) and target replacement (p < 0.001) are heavily mobile elements associated (orange & blue). Gene function specialization was defined as a “generalizability score” (Purple bar; right Y-axis). An inverse relationship was observed between how mobile a resistance mechanism is (left Y-axis) and its resistance generalizability. The least mobile group, reduced permeability, has the highest generalizability (9.5) while the mobile groups (Target Protection, Target Inactivation, and Target Replacement) have the lowest generalizability (1-2).

Genes mediating reduced permeability, efflux, and target alteration were predominantly located in non-GI chromosomal regions (p < 0.001), whereas target protection, inactivation, and replacement genes were strongly associated with mobile elements (p < 0.001). To quantify functional specialization, a “generalizability score” was calculated as the average number of drug classes to which each gene confers resistance. An inverse relationship was observed between mobility and generalizability. Mobile AMR genes (target replacement, inactivation, protection) had low generalizability scores (1–2), while genes in non-GI regions exhibited higher scores. For instance, genes reducing permeability had the fewest mobile determinants but the highest generalizability score (9.52; Fig. 3).

**Figure 3.**
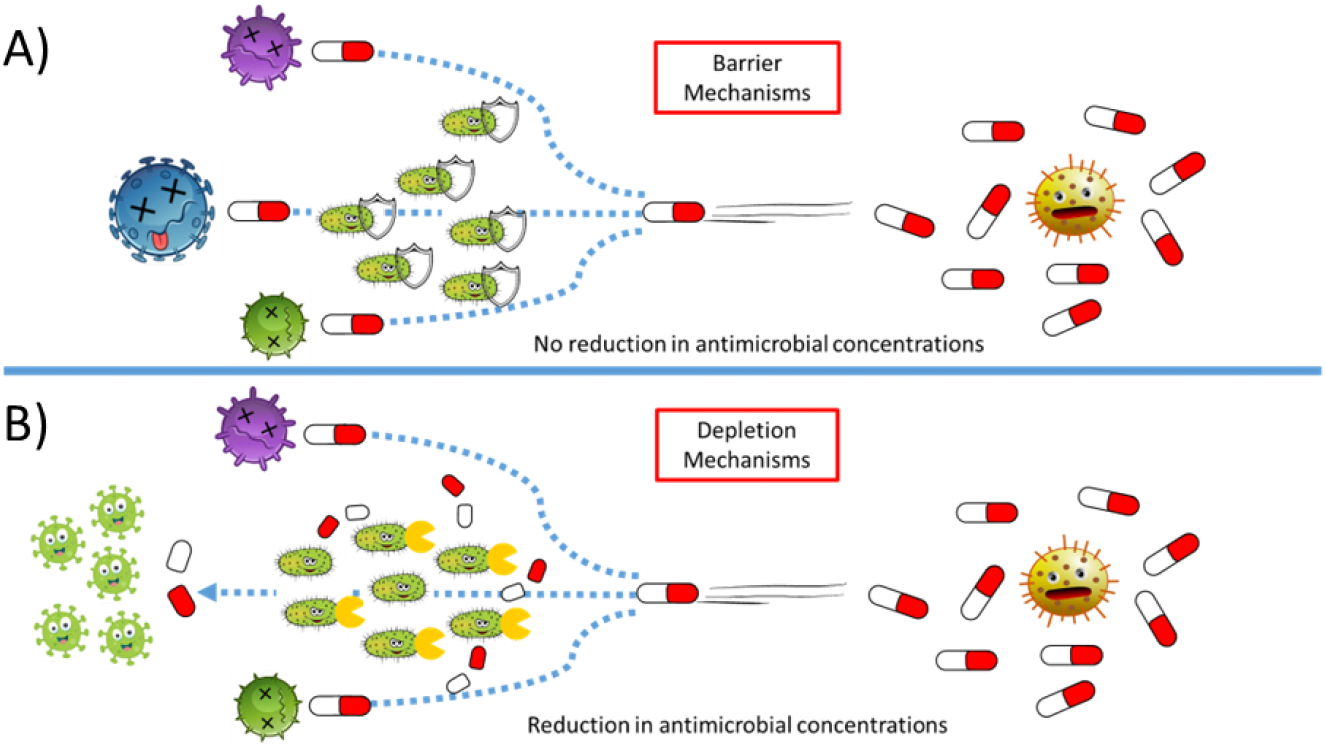
The effect of Barrier versus Depletion resistance mechanisms. A) Barrier mechanisms confer antimicrobial resistance via a passive blocking or removal of the antibiotic from the cell that does not affect environmental concentration. Therefore, the antimicrobials are capable of killing other bacterial species. This category includes Reduced Permeability to Antibiotic, Antibiotic Efflux, Antibiotic Target Alteration, and Antibiotic Target Protection. B) Depletion mechanisms confer active degradation or sequestration of antibiotics of secreted enzymes, removing it from the environment. Thus, the latter can allow the survival of other bacterial species in a cooperative environment. Depletion methods include Antibiotic Inactivation and Antibiotic Target Replacement, and are disproportionately encoded in mobile sequence regions, consistent with the “ecologically beneficial shared goods” that they represent.

#### AMR gene mobility patterns are confirmed in reduced subsets of the data

To further confirm these trends, we analyzed subsets corresponding to five WHO priority pathogen genera (*Pseudomonas, Salmonella, Klebsiella, Escherichia*, and *Burkholderia*). Across all subsets, AMR genes remained disproportionately associated with mobile elements. Resistance mechanism and generalizability consistently emerged as the primary determinants of AMR gene mobility.

### 6.3 Discussion

We conducted a large-scale analysis of AMR gene mobility across high-quality reference genomes and confirmed strong associations between known AMR determinants and the mobile genetic pool, collectively for a dataset of thousands of diverse bacterial species. Associations were unaffected by analyzing a dataset with reduced sampling bias. Associations were generally not affected by bacterial biological and genomic traits but depended on the AMR gene itself. Specifically, the gene’s underlying resistance mechanism and its generalizability were the primary determinants of mobility. These patterns persisted even when examining subsets of bacterial families, suggesting that AMR mobility is largely dictated by gene function rather than the host bacterium.

The Antibiotic Resistance Ontology (ARO), a hierarchical controlled vocabulary linking antimicrobial molecules, targets, resistance mechanisms, genes, and mutations (16), enabled us to explore the functional determinants of AMR mobility. Certain clinically relevant drug classes (e.g., carbapenemases that fall within antibiotic inactivation) were disproportionately associated with mobile elements, consistent with their rapid dissemination in nosocomial infections. Resistance mechanisms such as reduced permeability, efflux, and target alteration were predominantly chromosomally encoded, whereas target protection, inactivation, and target replacement were strongly associated with mobile elements. To conceptualize these trends, we categorized mechanisms into barrier mechanisms, which confer resistance by limiting intracellular antibiotic without altering environmental concentrations (reduced permeability, efflux, target alteration, target protection), and depletion mechanisms, which actively remove or neutralize antibiotics in the environment (inactivation, target replacement) (Fig. 3). Barrier mechanisms primarily benefit the individual bacterial cell or population, whereas depletion mechanisms may provide a shared advantage that benefits the community and potentially promotes horizontal gene transfer. The exception to this model was target protection, largely represented by Qnr proteins, which are mobile despite being to some degree a barrier mechanism. However, Qnr fluoroquinolone resistance genes emerged very recently and illustrate that AMR mobility can be influenced not only by mechanism but also by temporal factors: more recently acquired genes are more likely to reside on mobile elements.

Our analysis also indicates that specialized AMR genes are disproportionately mobile. Specialized functions, such as beta-lactamase-mediated antibiotic inactivation, are typically drug-specific and impose higher fitness costs compared with generalized mechanisms, like efflux pumps (24, 25). Mobility of these genes may mitigate host fitness costs or facilitate community-level interactions. Analogous to virulence factors and their association with mobile sequences (26), horizontal transfer of high-cost AMR genes may prevent “cheater” strains from outcompeting cooperative populations, consistent with the ecological public goods theory (27). Mobile specialized genes can distribute resistance across a community without requiring all members to carry the gene, increasing overall population fitness.

Subtle associations with bacterial traits were observed. Excluding plasmid-borne genes, AMR genes were more frequently localized to genomic islands in Gram-positive than Gram-negative species. This difference may reflect differences in cell envelope architecture. In Gram-negative diderms, plasmids can be efficiently transferred via conjugative pili, facilitating AMR gene movement across the diderm membranes. Consistent with this, Gram-negatives showed fewer GI-associated AMR genes and a greater reliance on plasmid-mediated resistance compared to Gram-positive species. Ecological isolation may also shape AMR gene distribution. As intracellular bacteria with limited horizontal gene transfer (e.g. *Mycobacterium)* disproportionately retained AMR genes on non-GI chromosomes because resistance tends to arise from substitutions in drug target proteins (28), whereas species naturally competent for DNA uptake like *Neisseria* species accessed mobile genetic pools readily. Nevertheless, resistance mechanism and generalizability remained the dominant determinants of mobility detected in our analysis.

Several limitations for this study should be considered. Existing literature disproportionately centers on high-consequence human- and animal-associated species, and thus naturally biasing toward more mobile AMR genes (e.g., carbapenemases). In turn, the RefSeq dataset overrepresents these hosts, and the CARD database is biased toward pathogen associated AMR determinants. While we analyzed reduced datasets to in part address this, our analysis does not fully eliminate these biases. Genomic island prediction using IslandViewer 4 also has high precision but may underpredict mobile regions and will not predict very small mobile genetic elements (21). However, based on past studies (29, 30, 31), AMR genes in the context of genomic islands are considered to be more clinically relevant, and reside on well-characterized islands of sufficient size for prediction. It should also be appreciated that analyses based on features like Gram status are computationally predicted, there may be errors for newly discovered environmental species. We also note that AMR mobility may be shaped by principles related to the complexity hypothesis, whereby genes encoding products with fewer functional interactions are more readily horizontally transferred than highly interconnected genes (32). Not all AMR genes act or move independently, as interacting resistance determinants are frequently co-localized on the same mobile genetic elements and therefore co-transferred, complicating attribution of mobility to individual gene properties (33, 34, 35). Consequently, observed associations between resistance mechanisms and mobility may partly reflect physical linkage and shared transfer histories of gene clusters rather than direct selective advantages of single genes that warrants further investigation. Despite these limitations, the observed trends did support more comprehensively, based on multiple large-scale datasets, that known AMR genes are disproportionately associated with mobile elements, particularly those with more specialized functions. This has important implications for AMR risk assessment, suggesting, for example, that antimicrobials with highly specific associated AMR mechanisms are at higher risk of losing effectiveness. Approaches for antimicrobial stewardship should factor in more the mechanisms of AMR into forecasting models, and development of approaches to combat AMR.

In conclusion, AMR mobility is influenced by multiple interacting factors, including fitness cost, selective pressure, temporal emergence, connectivity with other AMR and non-AMR proteins, and whether they can act as ecological public goods. Future work integrating improved predictions, more environmental sampling factoring in different niches and geography, and incorporating more experimental validation, could enable more quantitative modeling of AMR transmission. Understanding these patterns is critical for enhancing AMR surveillance and risk assessment across public health and agri-food systems, and for informing strategies to mitigate inappropriate antimicrobial use.

## 7. Figures and tables

**Supp Fig 1.**
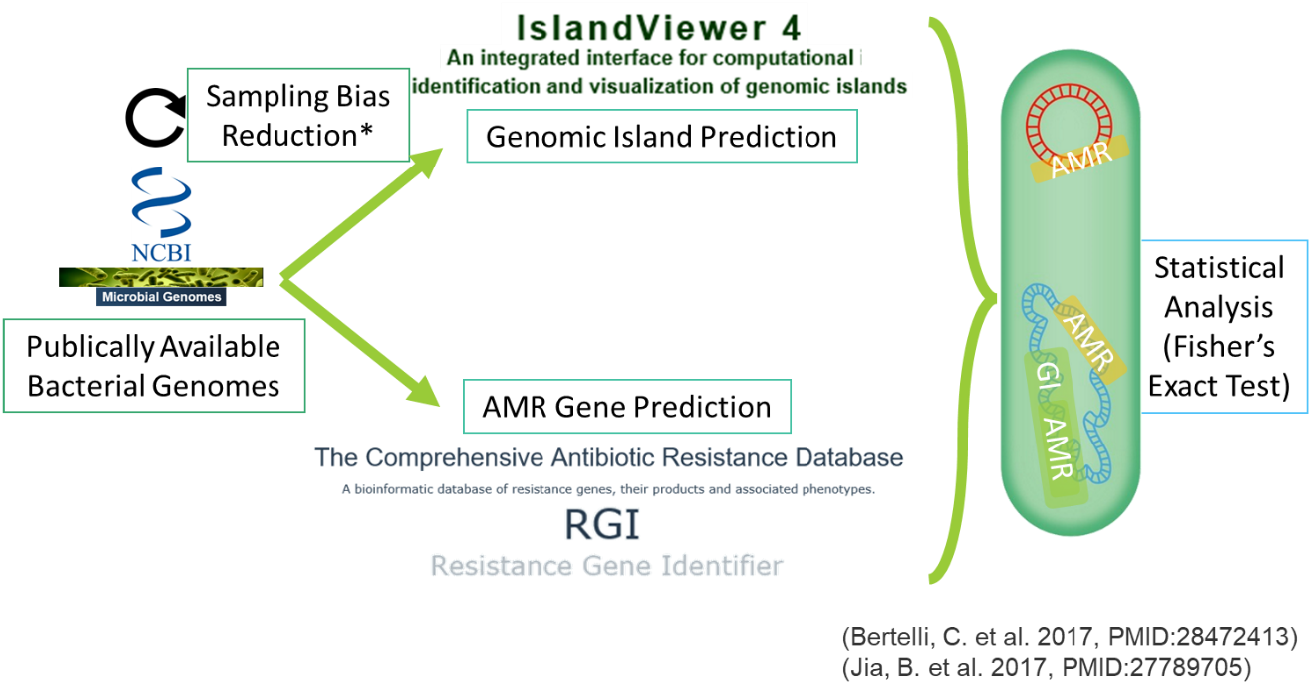
Overall workflow for understanding antimicrobial resistance gene mobility in RefSeq dataset. To explore and analyze the mobility of AMR genes, up to 300 bacterial chromosome and plasmid accessions from each genus present in the NCBI RefSeq database was randomly chosen to reduce the overrepresentation of certain species. GIs were then predicted using IslandViewer4 and AMR genes annotated with the Resistance Gene Identifier. Then combining the two prediction results, association of AMR genes within GIs, non-GI chromosomes and plasmids were statistically tested using Fisher’s Exact Test.

**Supp Fig 2.**
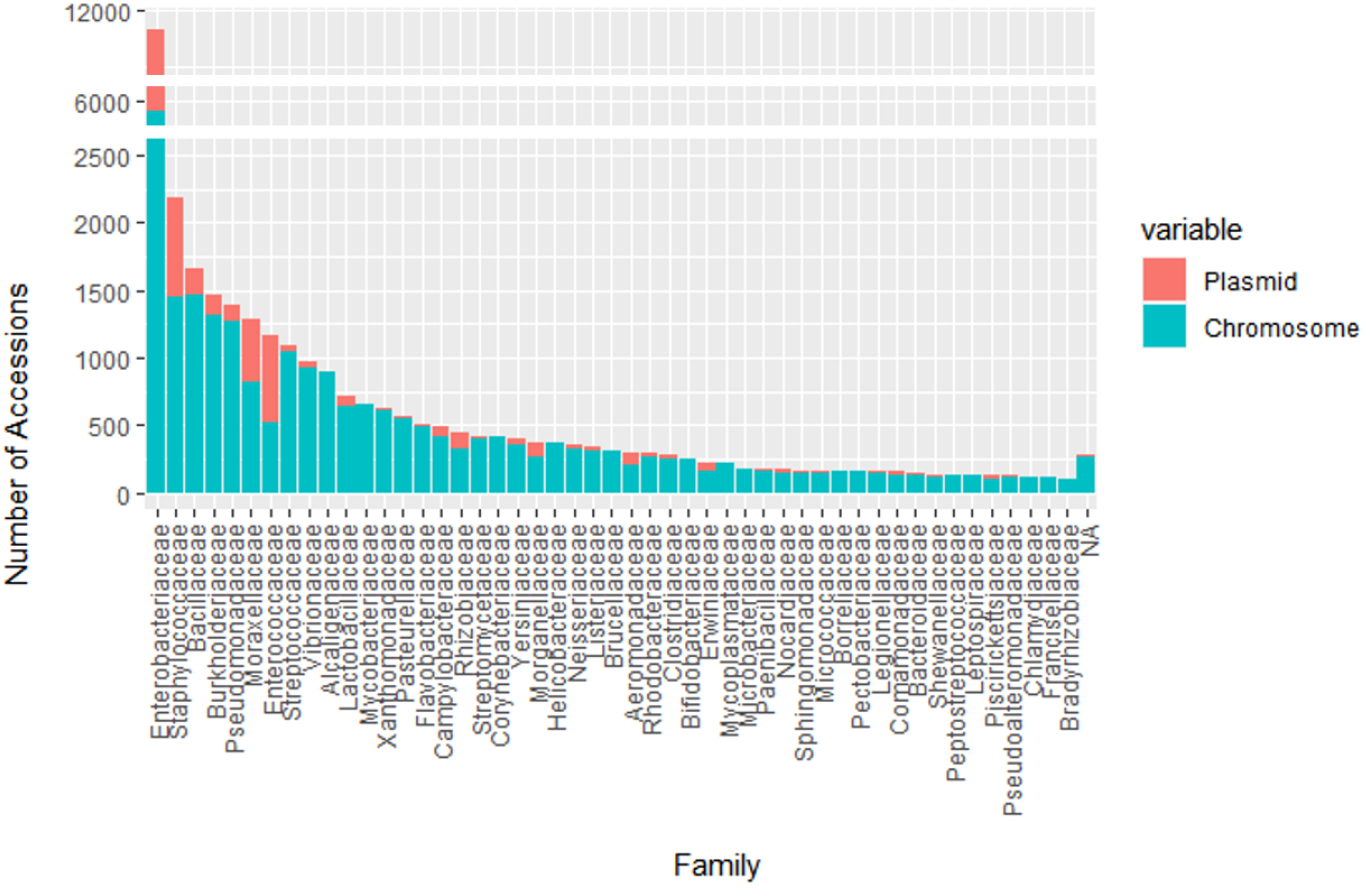
Summary of accessions in the RefSeq by the bacterial Family. The RefSeq bacterial dataset is overrepresented by species of human and animal relevance, namely pathogens as they are more commonly sequenced to date. The family Enterobacteraceae represents over 30% of all accessions (∼11,500) in the dataset with the top 10 family making up >50% of all accessions with an exponential decrease across the family.

**Supp Fig 3.**
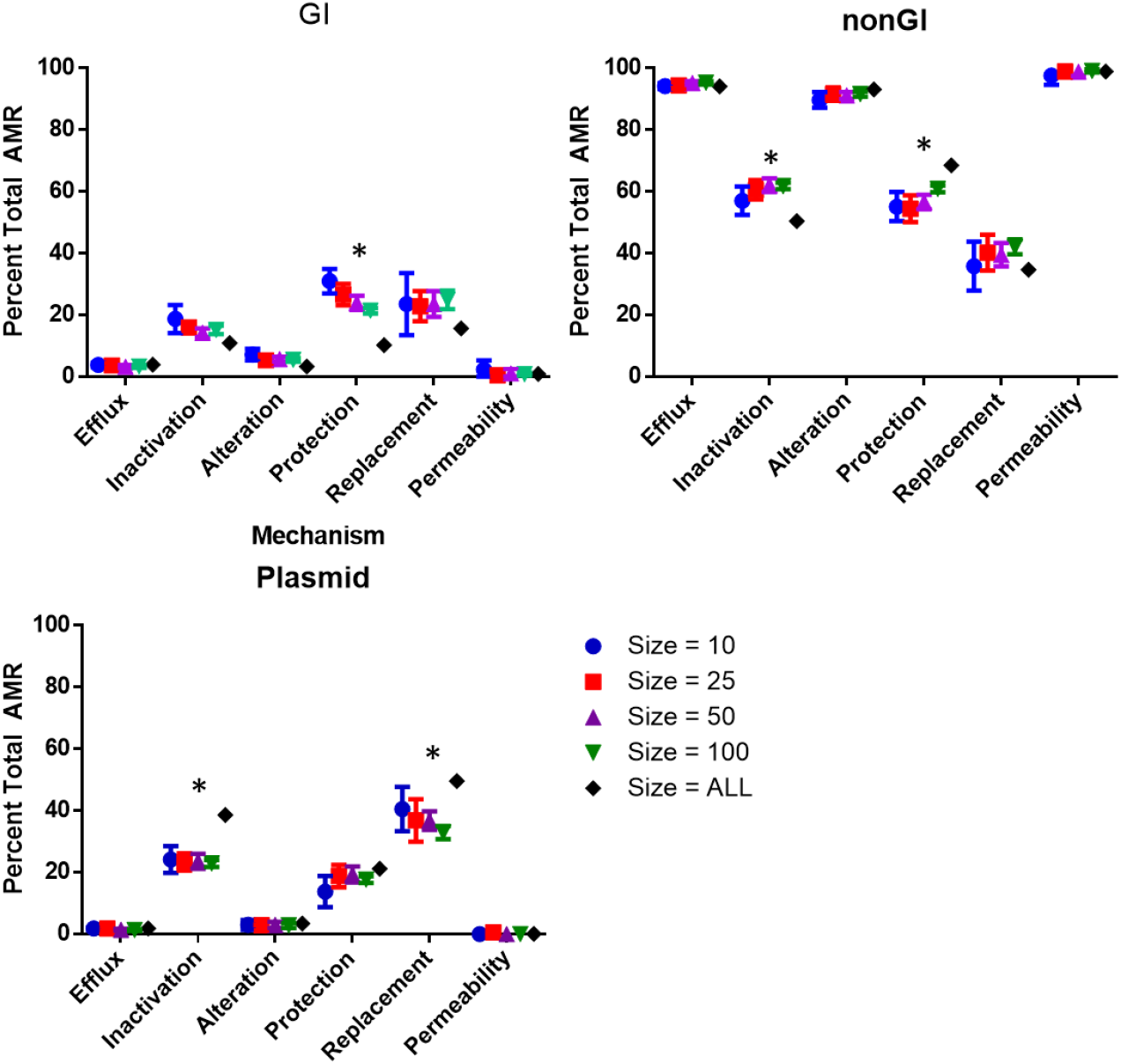
Limiting the number of accessions per genus changed the relative proportions of certain AMR gene mechanism classes, while the overall association with mobile elements remained unchanged. AMR gene resistance mechanisms were used as a proxy to assess changes across different amounts of downsampling. No significant differences were observed among datasets with 10 to 100 accessions per genus (circle, square, up arrow, down arrow). However, compared to the full dataset (diamond), there was a significant decrease in target protection associated resistance genes on genomic islands (GI), counterbalanced by an increased frequency on non-GI chromosomes (nonGI). Similarly, the full RefSeq dataset exhibited an increased frequency of antibiotic inactivation associated resistance genes on plasmids, with a corresponding decrease on non-GI chromosomes. These differences likely reflect the overrepresentation of Enterobacteriaceae in RefSeq.

**Supplementary Figure 4.**
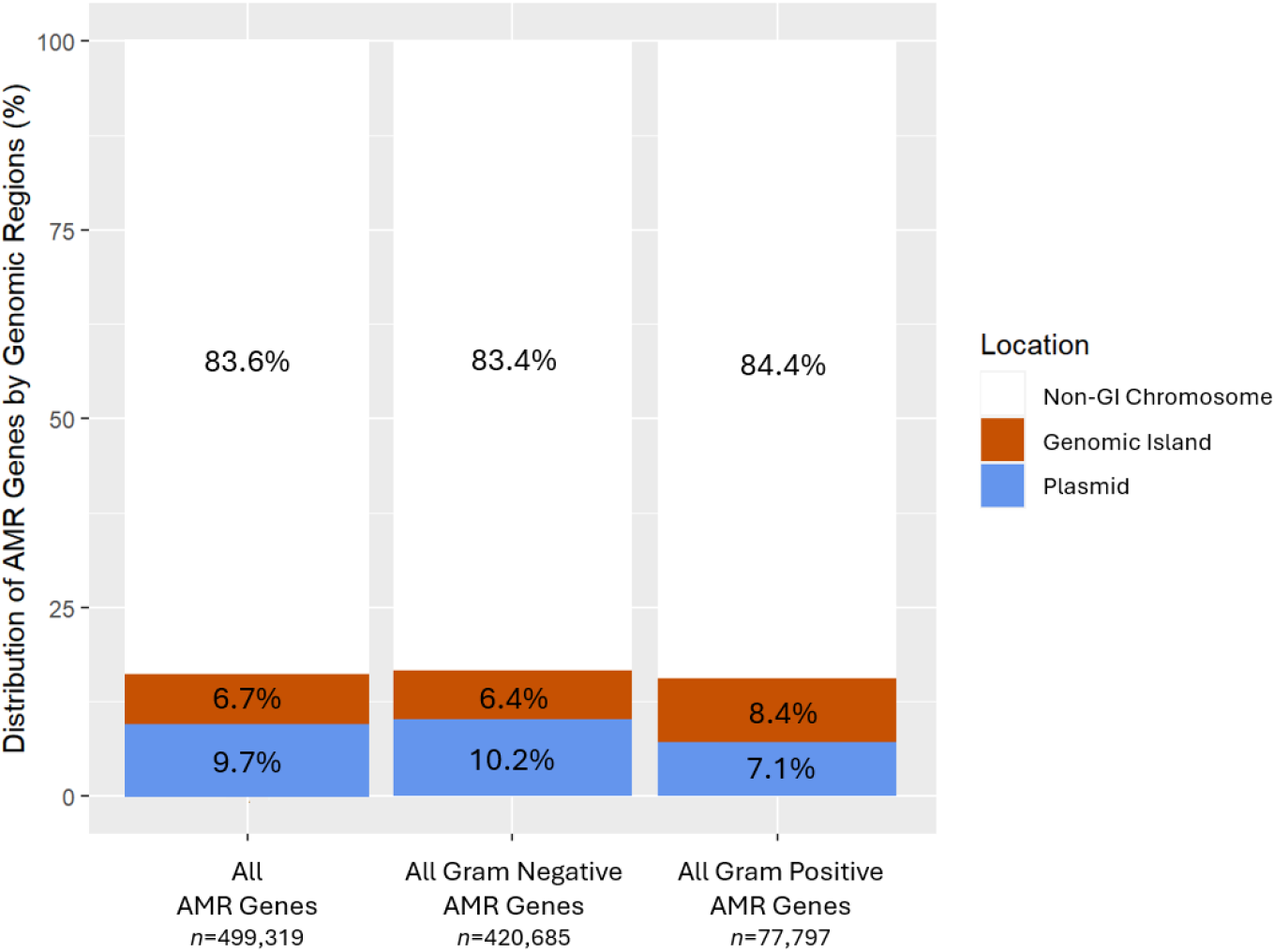
Predicted Gram status of the bacterial species did not impact the association between AMR genes and mobile genetic elements. When breaking down the AMR Genes by predicted Gram status, AMR genes in both Gram-positive and Gram-negative species were disproportionately found on mobile genetic elements (Fisher’s test, p < 0.001) similar to all AMR genes. A subtle difference is that Gram-negative bacteria had a higher proportion of AMR genes being present on mobile genetic elements, including both genomic islands (blue) and plasmid (red) compared to Gram-positive bacteria. However, the mobile AMR genes are disproportionally associated with GIs than plasmids in Gram-positives vs Gram-negatives. AMR Genes on genomes with atypical or indeterminate Gram status are not shown.

## 8 Author statements

### 8.1 Author contributions

CRediT: Conceptualization: **BJ, CB, FSLB**; Data curation: **BJ, JS, BPA**; Formal Analysis: **BJ, RGB, AGM, FSLB;** Funding acquisition: **RGB, AGM, FSLB**; Investigation: **BJ, CB**; Methodology: **BJ, BPA, ARR, RGB, FM, AGM, CB, FSLB;** Project administration: **RGB, FM, AGM, FSLB;** Resources, **BPA, ARR, AGM, FSLB**; Software: **BJ**; Supervision: **CB, FSLB**; Validation: **BJ**; Visualization: **BJ**; Writing – original draft: **BJ, FSLB;** Writing – review & editing: **All**

### 8.4 Ethical approval

Not Required

### 8.5 Consent for publication

Not Required

### 8.6 Acknowledgements

AGM was supported by a Cisco Research Chair in Bioinformatics (Cisco Systems Canada, Inc.) and a David Braley Chair in Computational Biology, generously supported by the family of the late Mr. David Braley. Fiona Brinkman holds a Simon Fraser University Distinguished Professorship. We acknowledge all the researchers who have provided data openly and freely to the database resources we used.

## References

1. Smith RA, M’ikanatha NM, Read AF. Antibiotic resistance: a primer and call to action. Health Commun. 2015;30(3):309–14. doi:10.1080/10410236.2014.943634.

2. Abbas A, Barkhouse A, Hackenberger D, Wright GD. Antibiotic resistance: a key microbial survival mechanism that threatens public health. Cell Host Microbe. 2024;32(6):837–51. doi:10.1016/j.chom.2024.05.015.

3. Naghavi M, Vollset SE, Ikuta KS, Swetschinski LR, Gray AP, Wool EE, et al. Global burden of bacterial antimicrobial resistance 1990–2021: a systematic analysis with forecasts to 2050. Lancet. 2024;404(10459):1199–226. doi:10.1016/S0140-6736(24)01867-1.

4. O’Neill J. Antimicrobial resistance: tackling a crisis for the future health and wealth of nations. London: Review on Antimicrobial Resistance; 2014.

5. World Health Organization. Global antibiotic resistance surveillance report 2025: WHO Global Antimicrobial Resistance and Use Surveillance System (GLASS). Geneva: World Health Organization; 2025.

6. Barlow M, Hall BG. Phylogenetic analysis shows that the OXA β-lactamase genes have been on plasmids for millions of years. J Mol Evol. 2002;55(3):314–21. doi:10.1007/s00239-002-2328-y.

7. Hall BG, Barlow M. Evolution of the serine β-lactamases: past, present and future. Drug Resist Updat. 2004;7(2):111–23. doi:10.1016/j.drup.2004.02.003.

8. Hall BG, Salipante SJ, Barlow M. Independent origins of subgroup B1+B2 and subgroup B3 metallo-β-lactamases. J Mol Evol. 2004;59(1):133–41. doi:10.1007/s00239-003-2572-9.

9. von Wintersdorff CJH, Penders J, van Niekerk JM, et al. Dissemination of antimicrobial resistance in microbial ecosystems through horizontal gene transfer. Front Microbiol. 2016;7:173. doi:10.3389/fmicb.2016.00173.

10. D’Costa VM, King CE, Kalan L, et al. Antibiotic resistance is ancient. Nature. 2011;477(7365):457–61. doi:10.1038/nature10388.

11. Bhullar K, Waglechner N, Pawlowski A, et al. Antibiotic resistance is prevalent in an isolated cave microbiome. PLoS One. 2012;7(4):e34953. doi:10.1371/journal.pone.0034953.

12. Poirel L, Rodriguez-Martinez JM, Mammeri H, Liard A, Nordmann P. Origin of plasmid-mediated quinolone resistance determinant QnrA. Antimicrob Agents Chemother. 2005;49(8):3523–5. doi:10.1128/AAC.49.8.3523-3525.2005.

13. Rodriguez MM, Power P, Radice M, Vay C, Famiglietti A, et al. Chromosome-encoded CTX-M-3 from Kluyvera ascorbata: a possible origin of plasmid-borne CTX-M-1-derived cefotaximases. Antimicrob Agents Chemother. 2004;48(12):4895–7. doi:10.1128/AAC.48.12.4895-4897.2004.

14. Doublet B, Boyd D, Mulvey MR, Cloeckaert A. The Salmonella genomic island 1 is an integrative mobilizable element. Mol Microbiol. 2005;55(6):1911–24. doi:10.1111/j.1365-2958.2005.04520.x.

15. Pillonetto M, Arend L, Vespero EC, Pelisson M, Chagas TPG, Carvalho-Assef APD, et al. First report of NDM-1-producing Acinetobacter baumannii sequence type 25 in Brazil. Antimicrob Agents Chemother. 2014;58(12):7592–4. doi:10.1128/AAC.03444-14.

16. Alcock BP, Huynh W, Chalil R, Smith KW, Raphenya AR, Wlodarski MA, et al. CARD 2023: expanded curation, support for machine learning, and resistome prediction at the Comprehensive Antibiotic Resistance Database. Nucleic Acids Res. 2023;51(D1):D690–9. doi:10.1093/nar/gkac920.

17. Bertelli C, Tilley KE, Brinkman FSL. Microbial genomic island discovery, visualization and analysis. Brief Bioinform. 2019;20(5):1685–98. doi:10.1093/bib/bby042.

18. Langille MGI, Hsiao WWL, Brinkman FSL. Detecting genomic islands using bioinformatics approaches. Nat Rev Microbiol. 2010;8:373–82. doi:10.1038/nrmicro2350.

19. Langille MG, Laird MR, Hsiao WW, Chiu TA, Eisen JA, Brinkman FS. MicrobeDB: a locally maintainable database of microbial genomic sequences. Bioinformatics. 2012;28(14):1947–8. doi:10.1093/bioinformatics/bts273.

20. World Health Organization. WHO bacterial priority pathogens list, 2024. Geneva: World Health Organization; 2024.

21. Bertelli C, Laird MR, Williams KP, et al. IslandViewer 4: expanded prediction of genomic islands for larger-scale datasets. Nucleic Acids Res. 2017;45(W1):W30–5. doi:10.1093/nar/gkx343.

22. Lau WYV, Hoad GR, Jin V, Winsor GL, Madyan A, Gray KL, et al. PSORTdb 4.0: expanded and redesigned bacterial and archaeal protein subcellular localization database. Nucleic Acids Res. 2021;49(D1):D803–8. doi:10.1093/nar/gkaa1095.

23. Wickham H. ggplot2: elegant graphics for data analysis. New York: Springer; 2016.

24. Pacheco JO, Alvarez-Ortega C, Rico MA, Martínez JL. Metabolic compensation of fitness costs is a general outcome for antibiotic-resistant Pseudomonas aeruginosa mutants overexpressing efflux pumps. mBio. 2017;8(4):e00500–17. doi:10.1128/mBio.00500-17.

25. Marciano DC, Karkouti OY, Palzkill T. A fitness cost associated with the antibiotic resistance enzyme SME-1 β-lactamase. Genetics. 2007;176(4):2381–92. doi:10.1534/genetics.106.069443.

26. Sui, S. J. H., Fedynak, A., Hsiao, W. W. L., Langille, M. G. I., & Brinkman, F. S. L. (2009). The Association of Virulence Factors with Genomic Islands. PLoS ONE, 4(12), e8094. 10.1371/journal.pone.0008094

27. Olson M. The logic of collective action: public goods and the theory of groups. Cambridge (MA): Harvard University Press; 1971.

28. Nimmo C, Millard J, Faulkner V, Monteserin J, Pugh H, Johnson EO. Evolution of Mycobacterium tuberculosis drug resistance in the genomic era. Front Cell Infect Microbiol. 2022 Oct 7;12:954074. doi: 10.3389/fcimb.2022.954074. PMID: 36275027; PMCID: PMC9585206.

29. Zhang H, Wu T, Ruan H. Unveiling genomic islands hosting antibiotic resistance genes and virulence genes in foodborne multidrug-resistant pathogenic Proteus vulgaris. Biology (Basel). 2025;14(7):858. doi:10.3390/biology14070858.

30. Audrey B, Cellier N, White F, Jacques P, Burrus V. A systematic approach to classify and characterize genomic islands driven by conjugative mobility using protein signatures. Nucleic Acids Res. 2023;51(16):8402–12. doi:10.1093/nar/gkad644.

31. Munshi ID, Mathuria A, Sharma H, Acharya M, Chaudhary A, Jain K, et al. Emerging concept of genomic islands in bacterial adaptation and pathogenicity. Res Microbiol. 2025;176(7):104303. doi:10.1016/j.resmic.2025.104303.

32. Cohen O, Gophna U, Pupko T. The complexity hypothesis revisited: connectivity rather than function constitutes a barrier to horizontal gene transfer. Mol Biol Evol. 2011;28(4):1481–9. doi:10.1093/molbev/msq333.

33. Li X, Brejnrod A, Trivedi U, Russell J, Thorsen J, Shah SA, et al. Co-localization of antibiotic resistance genes is widespread in the infant gut microbiome and associates with an immature gut microbial composition. Microbiome. 2024;12(1):87. doi:10.1186/s40168-024-01800-5.

34. Baker-Austin C, Wright MS, Stepanauskas R, McArthur JV. Co-selection of antibiotic and metal resistance. Trends Microbiol. 2006;14(4):176–82. doi:10.1016/j.tim.2006.02.006.a

35. Cantón R, Ruiz-Garbajosa P. Co-resistance: an opportunity for bacteria and resistance genes. Curr Opin Pharmacol. 2011;11(5):477–85. doi:10.1016/j.coph.2011.07.007.

